# The Proteome as a Whole: From a Passive Reflection of the Transcriptome to an Active Cellular Organizer

**DOI:** 10.64898/2026.09.15.751782

**Authors:** Anastasia Lazareva, Daria Matyushkina, Elena Vasilyeva, Alexey Kovalenko, Ivan Butenko, Vadim Govorun

## Abstract

Despite advances in proteomics, the proteome is still often viewed as a passive reflection of the transcriptome. Using *Mycoplasma gallisepticum* as a minimal cell model, we analyzed large-scale datasets across multiple stress conditions to investigate whether the proteome functions as an integrated regulatory system. The proteome remained quantitatively stable under stress, yet the bacteria adapted successfully. We identified a minimal core of 17 major proteins constituting 35% of cellular protein mass. These proteins exhibit distinct physicochemical properties—higher positive charge, lower hydrophilicity, and enrichment in intrinsically disordered regions— suggesting they shape cytoplasmic organization and may drive liquid–liquid phase separation. Minor proteins form tightly correlated clusters, while stress conditions trigger rearrangement of protein complexes without abundance changes, representing an energy-efficient adaptation strategy. Cross linking mass spectrometry revealed condition specific interactome remodeling, which may underlie mycoplasma adaptation to diverse stresses. Our findings suggest that the proteome operates as an active cellular organizer, where physicochemical properties and dynamic complex rearrangements enable adaptation independently of transcriptional control. This framework may inform synthetic biology efforts to engineer minimal cells.

## Introduction

The first studies that could be classified as proteomics began in 1974 with the introduction of two-dimensional electrophoresis for mapping *Escherichia coli* proteins^1^. Since then, technological advances have transformed the field, making it possible to quantitatively profile entire cellular proteomes^2,3^ and even highly abundant proteins in single cells^4^.

Despite these advances, the proteome is still often viewed mainly as a downstream outcome of transcriptional changes, where each protein is assigned either an individual function or a role within functional complexes. Based on this view, the classical model of bacterial regulation assumes that cellular responses occur through sequential changes in transcription followed by translation. However, evidence from mycoplasmas suggests that this model is not universal. In these bacteria, which serve as a model of a minimal cell, proteome stability under stress conditions has been observed despite significant transcriptional changes^5,6^.

Moreover, regulation in living cells is achieved not only through direct genomic and transcriptomic control that subsequently influences translation, but also through a wide range of epigenetic mechanisms. In bacteria, epigenetic regulation includes DNA methylation, RNA modification, and chromatin-like structural changes mediated by nucleoid-associated proteins and small non-coding RNAs^7,8^.

In the genus *Mycoplasma*, epigenetic mechanisms appear to be considerably streamlined while remaining functionally important^9,10^. This streamlined regulation is likely linked to the reduced genome and the limited number of regulatory proteins compared to other bacteria^11^. At the same time, mycoplasmas display only minor protein changes under stress conditions, suggesting that epigenetic mechanisms may play a central role in cellular adaptation.

For example, transcriptional regulators such as MraZ employ a unique “cradle-like” DNA-binding motif to control essential cell division genes, illustrating how protein–DNA interactions can compensate for reduced regulatory complexity^12,13^. Likewise, the WhiA regulator in *M. gallisepticum,* which represses the *rpsJ* operon and senses ATP^14^, further connects the metabolic state of the cell to its regulatory machinery.

Notably, WhiA contains an Intrinsically Disordered Region (IDR) located between its two functional domains. This region provides structural flexibility and promotes multivalent interactions with various biomolecules, potentially facilitating liquid–liquid phase separation (LLPS) both *in vivo* and *in vitro*. Over the past decade, LLPS has emerged as a universal mechanism for the spatiotemporal organization of cellular processes, in which biomolecules assemble into dynamic droplet-like condensates that function as membraneless organelles. These condensates can reversibly form and dissolve in response to cellular signals and are involved in genome organization, cytoskeletal assembly, stress adaptation, and the development of neurodegenerative diseases^15,16,17,18^.

Furthermore, proteins in the cellular environment can be viewed as soft condensed matter formed not only due to high concentrations, but in particular due to weak intermolecular interactions and sensitivity to thermal fluctuations^19^. The free-energy landscapes governing protein conformational changes and protein–protein interactions contain multiple minima separated by low barriers, resulting in thermally activated transitions between different states^20^. This similarity in energy scales characterizes intracellular systems, where entropic effects and thermal fluctuations play central roles in determining structural and dynamic properties.

Biomolecular condensates formed through liquid–liquid phase separation of proteins and nucleic acids create distinct intracellular compartments with altered local concentrations and protein dynamics^21^. In addition, crowding effects within the cellular environment add another layer of complexity, as the high concentration of macromolecules (200–400 mg/mL) creates conditions analogous to concentrated colloidal suspensions^22^. Under these conditions, intermolecular interactions can stabilize protein complexes and alter binding equilibria, while hydrodynamic interactions influence the diffusion and mobility of individual proteins^23^.

Typically, these physical aspects are not considered in standard cellular protein inventories, although they undoubtedly influence cellular organization and may also contribute to adaptation under stress conditions. This is particularly interesting in the context of minimal cells such as mycoplasmas, whose size can vary substantially despite the relative stability of their proteome^24^.

Recently, it was demonstrated that soluble proteins are directed toward the advancing edge of the cell through advection, a process in which diffusion is enhanced by intracellular fluid flow^25^. This occurs within a specialized leading-edge compartment separated from the bulk cytoplasm by an actin–myosin condensate barrier. This mechanism synchronizes protein distribution with local changes in cell morphology and highlights the need for a broader view of the functional and regulatory roles of cellular components.

We hypothesize that, during the coordination of certain cellular processes, proteins may autoregulate independently of transcriptional control. Beyond their specific biological functions, proteins also possess physicochemical properties capable of influencing their surrounding microenvironment, much like any other molecules. The more abundant a protein is, the greater its potential physical impact, underscoring the importance of investigating quantitative patterns in proteins with distinct properties. In *E. coli*, proteome dynamics have been proposed as a mechanism for short-term “memory” of metabolic states^26,27^, suggesting that the proteome as a whole may possess broader functional roles, potentially even in more complex microorganisms.

Modern proteomic technologies allow highly accurate quantification of cellular composition. However, even a comprehensive protein inventory is insufficient to answer several fundamental physical and biochemical questions related to the functioning of a living cell. One particularly striking observation is that some proteins are present in cells at concentrations exceeding their solubility limits. Likewise, it remains unclear how molecular movement within the cell can occur at rates up to 1000 times faster than expected from diffusion alone^28^.

Another important concerns the degree of protein hydration and its influence on protein mobility and intermolecular interactions within the cell. Hydration plays a critical role in charge shielding and the regulation of molecular interactions. According to our calculations, in a bacterium such as *E. coli*, there is barely enough water to hydrate all cellular molecules. Approximately 40% of water molecules surround DNA and stabilize its structure^29^, while 55% to 70% are required to form protein hydration shells^30^. Similar estimates have been reported for human red blood cells, *Haloarcula marismortui*, *Bacillus subtilis*, and bacterial spores^31^.

These observations raise a fundamental question: what determines the physicochemical properties of the cellular interior and allows molecules to remain soluble above their water solubility limits? Theoretical studies suggest that proteins themselves may possess properties analogous to those of water molecules^32^. In addition, ATP, traditionally viewed as a universal energy carrier, has also been shown to perform a physicochemical hydrotropic function within the cell. In this context, ATP can be considered a small amphiphilic molecule that, at sufficiently high concentrations, increases the solubility of hydrophobic regions on biopolymers, thereby reducing their tendency toward unwanted aggregation^33^. ATP may also participate in phase separation together with proteins^34^.

Since one of the major goals of modern synthetic biology is the creation of an artificial cell through a bottom-up approach, understanding proteome organization in such systems has become a priority. Bacteria from the genus *Mycoplasma* are widely regarded as a classical model of a minimal cell. Decades of systematic research by multiple groups have generated extensive datasets describing the average abundance of individual cellular molecules across a broad range of conditions and physiological states.

These data not only enable the construction of numerical models for predicting cellular viability and functionality^35,36^, but also provide an opportunity to identify previously unrecognized organizational principles through large-scale meta-analysis—an approach undertaken in the present work. Using existing algorithms for predicting protein physicochemical properties together with accumulated large-scale datasets, we characterized each protein of the *Mycoplasma gallisepticum* proteome both qualitatively and quantitatively. We then extrapolated these data across the entire proteome to obtain a holistic view of the distribution of protein properties within the cell.

By analyzing the complete dataset, we identified subtle quantitative relationships among proteins that are characteristic of the proteome, both across all studied conditions collectively and within individual stress conditions. We identified and characterized a minimal core of proteins constituting the major fraction of the *M. gallisepticum* cellular proteome. In contrast, the remaining lower-abundance proteins showed a tendency to form highly correlated clusters.

Given the quantitative stability of the proteome under stress conditions, mycoplasmas likely respond through qualitative rearrangements of protein complexes (“protein collisions”), potentially enabling adaptation to new stressors without substantial energy expenditure. Collectively, these findings may indirectly indicate the existence of additional layers of proteomic regulation within the cell and highlight the importance of considering proteome dynamics as an integrated system rather than as isolated protein inventories.

## Results

### The *M. gallisepticum* Proteome Is Stable Under Any Conditions

Over years of systematic study of *M. gallisepticum* in our laboratory, we prepared and analyzed several thousand samples covering total cellular proteomes and various subcellular fractions under diverse conditions. This extensive dataset motivated us to pursue a comprehensive characterization of the mycoplasma proteome. Given that the mycoplasma proteome remains largely unchanged under various stress conditions while the cells retain remarkable adaptive capacity, we sought to investigate the types of interactions that occur within the proteome when a direct correlation with transcription is absent^5^.

To assess proteome behavior as an integrated system, we selected 504 samples representing the total proteome of *Mycoplasma gallisepticum* S6 from 30 independent experiments, including biological replicates. All samples were prepared using a standardized protocol and analyzed under identical conditions on the same Orbitrap Exploris 480 instrument. The analysis included cells grown under normal laboratory conditions (control) as well as under stress conditions, including acute heat shock (46°C for 30 min) and chronic hypo- and hyperosmotic stress. For osmotic stress experiments, cells were cultivated in media containing different NaCl concentrations, which differentially affected culture growth rates.

Our previous studies showed that at NaCl concentrations ranging from 30 mM to 160 mM, the metabolic rate did not differ significantly from that observed under control conditions (85 mM NaCl). In contrast, at 0 mM and 250 mM NaCl, the metabolic rate decreased approximately twofold^24^.

All samples were initially evaluated for technical quality, including missed cleavage rates, artifactual modifications, and contaminant protein levels. As shown in Figures 1a,b, the accumulated dataset exhibited low levels of missed cleavages and peptide side modifications, minimal contamination for bacterial proteomics samples, and high similarity between protein identifications across different conditions (p-value < 0.01).

**Figure 1.**
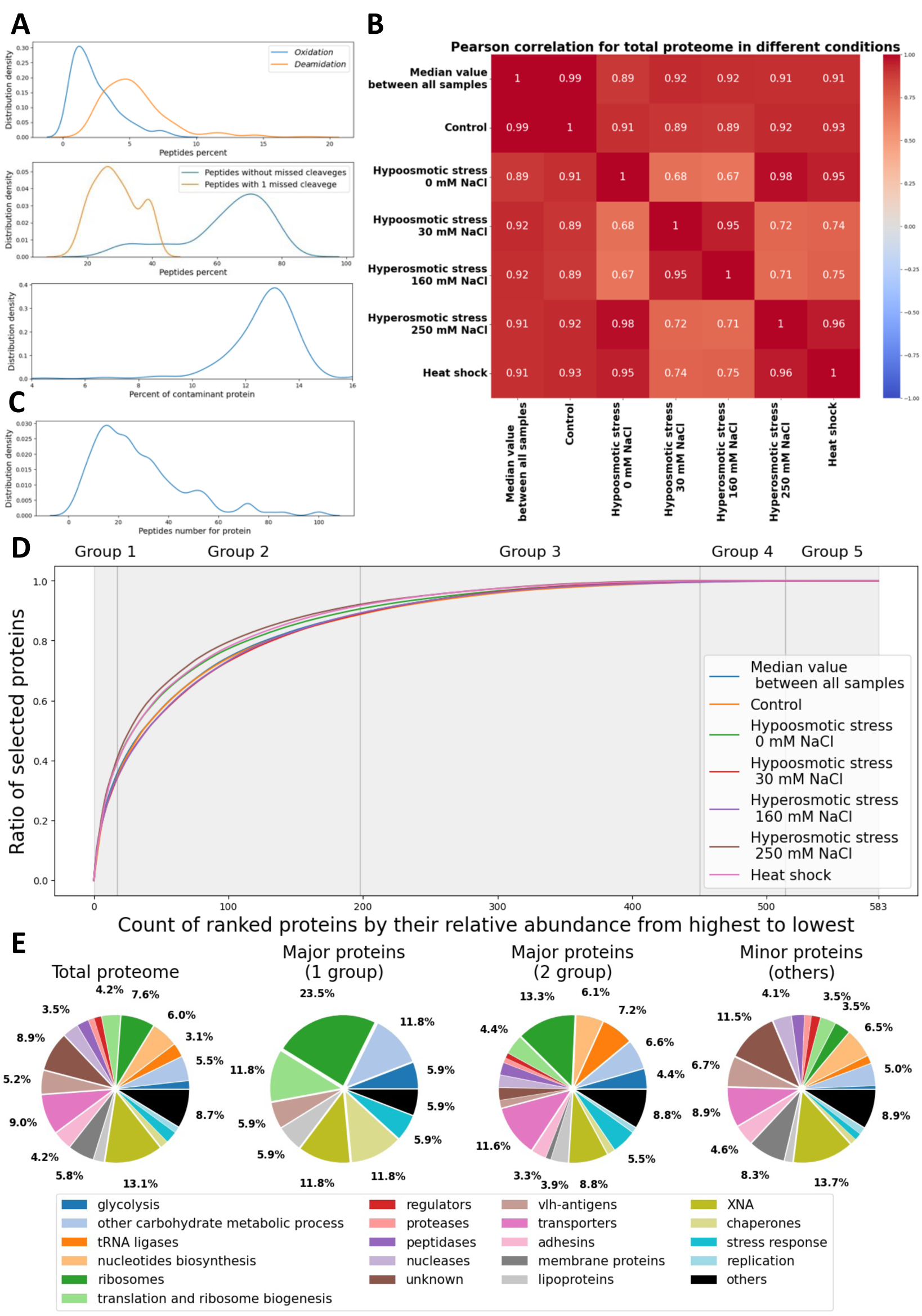
Proteomics characterization of *M. gallisepticum*. (A) Distribution of modified peptides, peptides with missed trypsin cleavages, and contaminant proteins across samples. (B) Heatmap showing Pearson correlation coefficients for collective protein abundance changes across conditions. (C) Distribution of the number of peptides identified per protein. (D) Proportion of the cellular proteome occupied by different numbers of proteins ranked by relative abundance from highest to lowest. (E) Functional classification of major proteins, minor proteins, and the total proteome according to biological function.

In total, proteins identified from the target organism covered approximately 85% of the annotated *Mycoplasma* proteome, corresponding to 688 proteins out of 812 annotated open reading frames. The remaining 124 proteins likely escaped detection due to their specific characteristics. Most of these were small proteins (up to 100 amino acids) with uncharacterized functions. Others are known to be present at low abundance, such as the Vlh antigens GCW_01910, GCW_02610, and GCW_02425^37,38^ or possess membrane localization, which complicates their extraction together with bulk proteins and may lead to losses during sample preparation.

We chose to examine all observed minor fluctuations in the *M. gallisepticum* proteome as part of a unified system in which all elements may be interconnected, contributing not only through their biological functions but also through their physicochemical properties. Since this type of proteome analysis is relatively novel and potentially susceptible to artifacts, the selection of high-quality and reliable data was particularly important. Consequently, we deliberately excluded proteins identified by only 1–2 peptides (Figure 1c), corresponding to 105 proteins.

After obtaining protein abundance values as iBAQ intensities under different conditions, we normalized each sample to the total abundance of all proteins within that sample. Median values were then calculated for each condition and subsequently renormalized to a total sum of 1, allowing more accurate comparisons on a common scale.

To assess proteome saturation, we ranked the remaining proteins (583 proteins, corresponding to 72% of the proteome) according to their relative abundance from highest to lowest (Figure 1d). Analysis of global proteome changes revealed that, under all stress conditions, the diversity of proteins constituting the majority of cellular mass decreased, resulting in a leftward shift of the curve, where fewer proteins accounted for the same fraction of total mass. However, changes in the abundance of major proteins did not exceed a twofold difference. Targeted analysis of these proteins (Supplementary Table 1) did not reveal functional enrichment relative to the proteome as a whole.

When proteins are considered not in terms of their biological functions, but rather through their physicochemical properties—such as charge, hydrophobicity, polarity, and reactivity—their fractional contribution to the total proteome mass becomes especially important. To evaluate the physical and functional roles of proteins at the proteome level, we stratified them into groups based on their average abundance across all studied conditions.

Our dataset contains relative protein abundances, where each major protein represents several hundredths of the total cellular protein mass. Proteins accounting for at least 0.01 of the normalized median value under control conditions were assigned to the first (major) order (Group 1). Proteins within the range [0.001;0.01) were assigned to the second order (Group 2), those within [0.0001;0.001) to the third order, and so forth (Table 1).

**Table 1.** General characteristics of *M. gallisepticum* protein groups based their quantitative representation.

|  | Total relative protein content | Number of proteins |
| --- | --- | --- |
| **Group 1**<br>**(>10<sup>-2</sup>)** | 0.353 | 17 |
| **Group 2**<br>($10^{-2} \gg 10^{-3}$) | 0.538 | 181 |
| **Group 3**<br>($10^{-3} \gg 10^{-4}$) | 0.105 | 252 |
| **Group 4**<br>($10^{-4} \gg 10^{-5}$) | 0.003 | 64 |
| **Group 5**<br>($10^{-5} \gg 10^{-6}$) | 0.0 | 69 |

Since only the first two groups contribute substantially to the total cellular protein mass (89%), subsequent analyses focused primarily on them. We first compared their qualitative composition using Gene Ontology terms. Major proteins from Groups 1 and 2 did not differ from lower-abundance proteins in the diversity of represented protein categories. However, they differed in the overall diversity levels of specific metabolic pathways (Figure 1e).

### Major Proteins May Define the Physicochemical Properties of the Entire Cytoplasm

For all *M. gallisepticum* proteins, we compiled a database of physicochemical characteristics (Supplementary Table 1). Protein length and molecular mass were obtained from UniProt. The ProtParam module of the Expasy server^39^ was used to calculate isoelectric point, total numbers of positively and negatively charged amino acids, instability index, aliphatic index, and GRAVY (Grand Average of Hydropathy).

Protein volume and packing density were calculated using a Java-implemented method^40^ based on structures predicted by AlphaFold^41^. Intrinsically disordered regions (IDRs) were predicted using the IUPred2A program^42^. Reactogenicity, defined here as the ability of a protein to interact with other proteins, was predicted using the ISPRED-SEQ software^43^. Protein solubility was predicted using the CamSol method^44^, while dipole moments were calculated using the ProteinDipole script^45^. Functional stability and melting temperatures were predicted using TemStaPro^46^ and DeepSTABp^47^ respectively. We assessed the connectivity of the obtained property data by computing Pearson correlation coefficients and evaluating their statistical significance. No significant association was observed between most of the examined properties, with two notable exceptions. First, the Aliphatic index and GRAVY exhibited an indirect relationship, attributable to the hydrophobic character of aliphatic amino acid residues. Second, a set of properties comprising protein length, total molecular volume, and the number of positively and negatively charged amino acids displayed interdependence that follows directly from their definitions, as these quantities are computed from the same underlying residue composition.

For major proteins, DNA-binding probability was estimated from AlphaFold-predicted structures using the DP-Bind web server^48^. Since DP-Bind generates predictions for individual amino acids, the fraction of amino acids predicted to bind DNA was used for subsequent analyses. We initially focused on Group 1, which consisted of 17 proteins, each contributing more than 1% to the total protein mass (Table 2). Since these proteins are functionally unrelated, we compared their physicochemical properties while accounting for their relative contribution to the overall proteome using weighted density distribution plots. Group 1 proteins displayed physicochemical characteristics distinct from those of the remaining proteome and may therefore serve as key drivers of local environmental changes and compartmentalization within specific cellular regions (Figure 2a).

**Figure 2.**
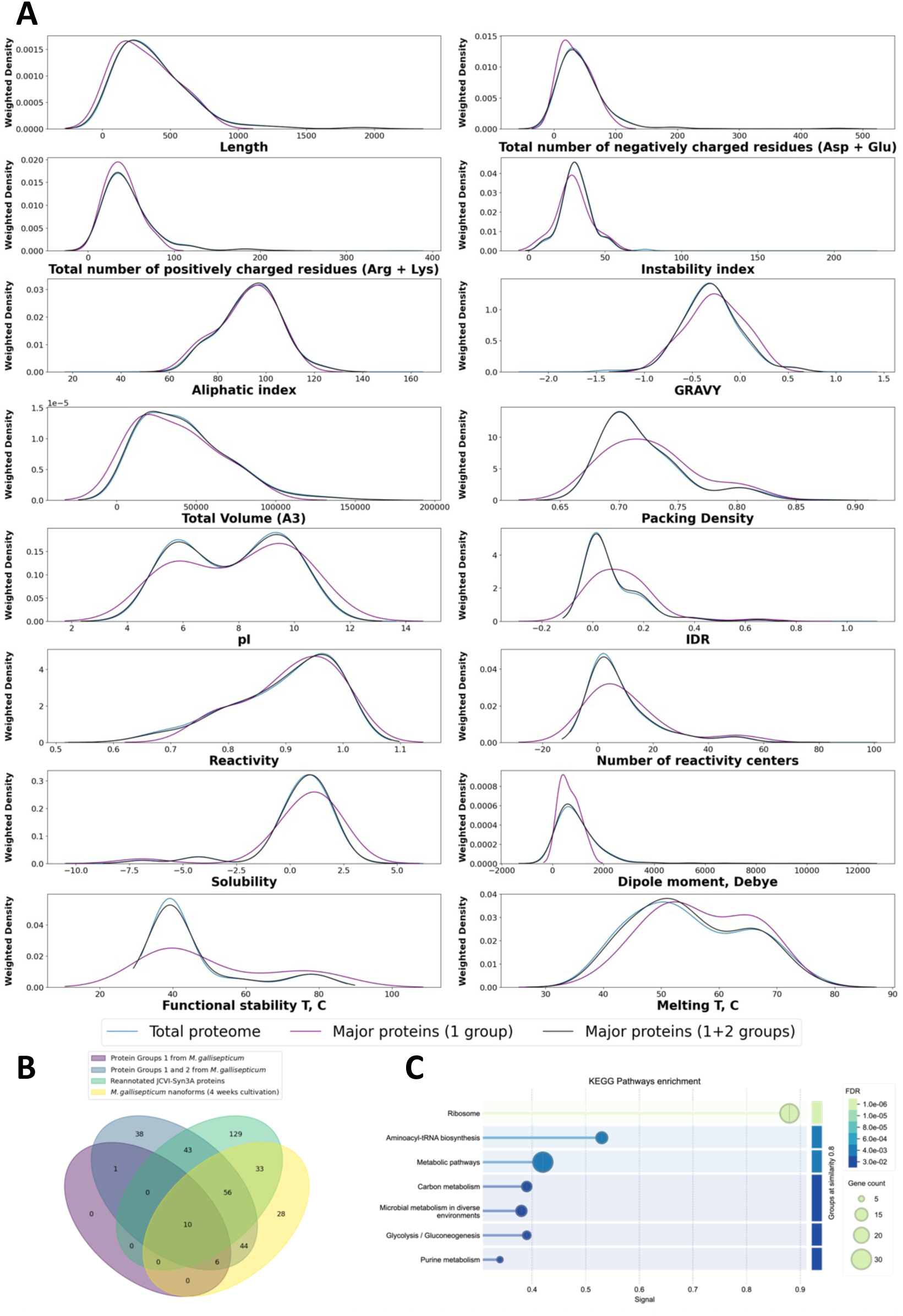
Property and functional characterization of *M. gallisepticum* proteins, which included in Groups 1 and 2. (A) Distributions of physicochemical properties for all proteins of the *M. gallisepticum* proteome, Group 1 proteins, and Groups 1+2 combined. (B) Venn diagram showing the overlap between Group 1 and Group 2 proteins from *M. gallisepticum* cells, proteins detected in *M. gallisepticum* nanoforms, and JCVI-Syn3A proteins^50^ re-annotated in *M. gallisepticum* terms. (C) Functional enrichment of Venn diagram intersection, comprising 109 proteins, performed using STRING-db^51^.

**Table 2.**
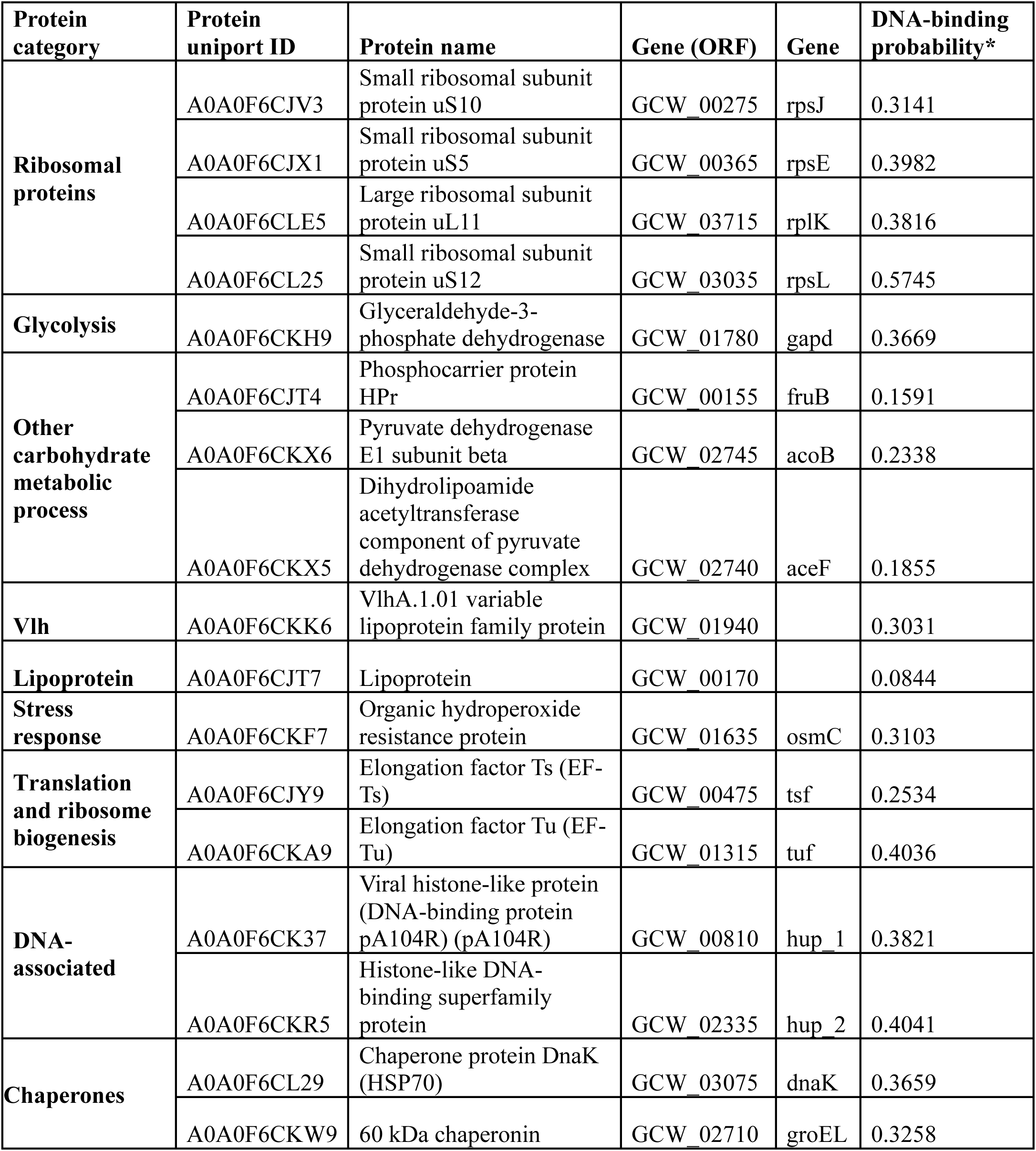
Major proteins of *M. gallisepticum* with a relative content ≥ 0.01.

| Protein category | Protein uniport ID | Protein name | Gene (ORF) | Gene | DNA-binding probability* |
| --- | --- | --- | --- | --- | --- |
| **Ribosomal proteins** | A0A0F6CJV3 | Small ribosomal subunit protein uS10 | GCW_00275 | rpsJ | 0.3141 |
|  | A0A0F6CJX1 | Small ribosomal subunit protein uS5 | GCW_00365 | rpsE | 0.3982 |
|  | A0A0F6CLE5 | Large ribosomal subunit protein uL11 | GCW_03715 | rplK | 0.3816 |
|  | A0A0F6CL25 | Small ribosomal subunit protein uS12 | GCW_03035 | rpsL | 0.5745 |
| **Glycolysis** | A0A0F6CKH9 | Glyceraldehyde-3-phosphate dehydrogenase | GCW_01780 | gapd | 0.3669 |
| **Other carbohydrate metabolic process** | A0A0F6CJT4 | Phosphocarrier protein HPr | GCW_00155 | fruB | 0.1591 |
|  | A0A0F6CKX6 | Pyruvate dehydrogenase E1 subunit beta | GCW_02745 | acoB | 0.2338 |
|  | A0A0F6CKX5 | Dihydrolipoamide acetyltransferase component of pyruvate dehydrogenase complex | GCW_02740 | aceF | 0.1855 |
| **Vlh** | A0A0F6CKK6 | VlhA.1.01 variable lipoprotein family protein | GCW_01940 |  | 0.3031 |
| **Lipoprotein** | A0A0F6CJT7 | Lipoprotein | GCW_00170 |  | 0.0844 |
| **Stress response** | A0A0F6CKF7 | Organic hydroperoxide resistance protein | GCW_01635 | osmC | 0.3103 |
| **Translation and ribosome biogenesis** | A0A0F6CJY9 | Elongation factor Ts (EF-Ts) | GCW_00475 | tsf | 0.2534 |
|  | A0A0F6CKA9 | Elongation factor Tu (EF-Tu) | GCW_01315 | tuf | 0.4036 |
| **DNA-associated** | A0A0F6CK37 | Viral histone-like protein (DNA-binding protein pA104R) (pA104R) | GCW_00810 | hup_1 | 0.3821 |
|  | A0A0F6CKR5 | Histone-like DNA-binding superfamily protein | GCW_02335 | hup_2 | 0.4041 |
| **Chaperones** | A0A0F6CL29 | Chaperone protein DnaK (HSP70) | GCW_03075 | dnaK | 0.3659 |
|  | A0A0F6CKW9 | 60 kDa chaperonin | GCW_02710 | groEL | 0.3258 |

Predicted using the DP-Bind web-server^48^. Values are generally considered indicative of DNA-binding propensity: the higher the value, the more likely the protein is to contain a complete DNA-binding domain.

On average, these proteins are characterized by a higher positive charge, largely due to the abundance of histone-like and ribosomal proteins, as well as by reduced hydrophilicity (lower GRAVY index). In addition, they contain more potential intrinsically disordered regions (IDRs) and reactogenicity sites, suggesting a possible scaffold role in biocondensate formation. We compared their sequences and structural organization with those of other mycoplasmas (*Mycoplasma genitalium* and *Mycoplasma pneumoniae*), including the minimal cell JCVI-Syn3A, as well as model organisms such as *Escherichia coli*, *Bacillus subtilis*, *Pseudomonas aeruginosa*, and *Staphylococcus aureus*, for proteins that have homologs in all of these organisms. We found that IDRs often display similar sequences and occupy similar regions within homologous proteins across these organisms (Supplementary Table 2).

Notably, major proteins are enriched not only in species with high melting points, but also in those that maintain functional activity at elevated temperatures. Several of these proteins are also classified as nucleic acid-binding. For most of them, with the exception of lipoprotein, FruB, and AceF, the predicted probability of DNA binding is of the same order of magnitude as that of established DNA-binding proteins (Table 2). Group 1 is significantly enriched in proteins that have both a high melting temperature and a high functional stability temperature, defined here as the temperature at which functional activity begins to decline. When Group 2 proteins are included, these features shift closer to the overall mycoplasma proteome profile, likely because these proteins account for the majority of total protein mass in the cell (Figure 2a).

### Major Cellular Proteins Constitute the Minimal Proteomic Core

Since Group 1 and Group 2 proteins together constitute 89% of the cellular protein biomass, essentially representing the major proteomic core, we compared them to existing minimal cell models. Mycoplasmas are known to form metastable nanoforms as an adaptation to prolonged non-culturable conditions^49^. These nanoforms are characterized by reduced metabolic activity, decreased cell size, lowered transcription levels, and diminished protein diversity. Notably, some nanoforms can revert to a normal state even after 4 weeks of incubation once returned to nutrient-rich conditions, suggesting that the nanoform proteome may itself represent a form of minimal proteome. Proteomic analysis of *M. gallisepticum* nanoforms, obtained after 4 weeks of cultivation in nutrient-depleted medium lacking lipids, amino acids, and nucleosides, confirmed that all Group 1 and Group 2 proteins were retained (Figure 2b).

Furthermore, the most well-known minimal cell, JCVI-Syn3A, a bacterium created by transplanting a chemically synthesized genome, was analyzed using published proteomic data^50^ to determine overlap between its proteomic composition and the identified “minimal core” of *M. gallisepticum* S6. Since these bacteria belong to different mycoplasma species, we performed a genomic comparison and found functional correspondence between 271 of the 455 JCVI-Syn3A genes with *M. gallisepticum* S6 genes.

A substantial fraction of the non-corresponding proteins lacked functional annotation, accounting for approximately 30% of the JCVI-Syn3A proteome (130 proteins). Similarly, 22% of the *M. gallisepticum* proteome (184 proteins) is also unannotated, which further limits the number of detectable matches between the two proteomes. The remaining proteins were insufficiently annotated in one or both organisms (Supplementary Table 3).

When the shared proteins were compared with our selected groups, Groups 1 and 2 from *M. gallisepticum* and 4-week nanoforms, 55% of the selected proteins were also found in JCVI-Syn3A (Figure 2b). Analysis of the intersecting proteins (Supplementary Table 3) showed enrichment for nearly all key metabolic pathways of mycoplasma^51^ (Figure 2c). Although most pathways were not fully represented, they retained core functional components, supporting the identification of this protein group as the cell’s “minimal core” or at least a substantial part of it.

### Adaptation To Stress Occurs Through The Rearrangement Of Complexes With Major Core Proteins

To better understand how the minimal protein core is organized within *M. gallisepticum*, we conducted an intracellular interactome analysis. For selected stress conditions, we performed a series of experiments using the cell-penetrating, flexible cross-linker EGS, which contains amine-reactive NHS-ester groups separated by a 16-atom spacer arm. EGS was added to grown and treated cultures to cross-link co-localized proteins, allowing interacting pairs to be identified by mass spectrometry.

The spacer length was selected to capture weak and transient protein interactions, rather than only stable canonical protein complexes. We then defined a “cross-linking frequency” metric as the proportion of biological replicates in which a given protein pair was detected. An interaction was considered reliable if it was detected in at least two out of three technical replicates within each of the seven biological replicates.

First, a global comparison of interaction occurrence frequencies was performed using the Pearson correlation coefficient (Figure 3a), revealing a lack of conserved interaction trends between proteins. We therefore focused specifically on interactions between Group 1 major proteins and all others proteins, regardless of their representation level. As shown by the UpSet plot (Figure 3b, Supplementary Table 4), only a small fraction of interactions was shared across all conditions. These included the interaction of RpsE with RpsB and AtpC, the interaction of Hup_2 with HisS and tRNA synthase TsaD; and interactions of the chaperone DnaK with six proteins: the methyltransferase RlmB, ribosomal protein RpsM, sigma factor GCW_00440, uncharacterized protein GCW_91851, putative ribosomal protein GCW_03230, and the M-domain IgG-blocking protein GCW_03985.

**Figure 3.**
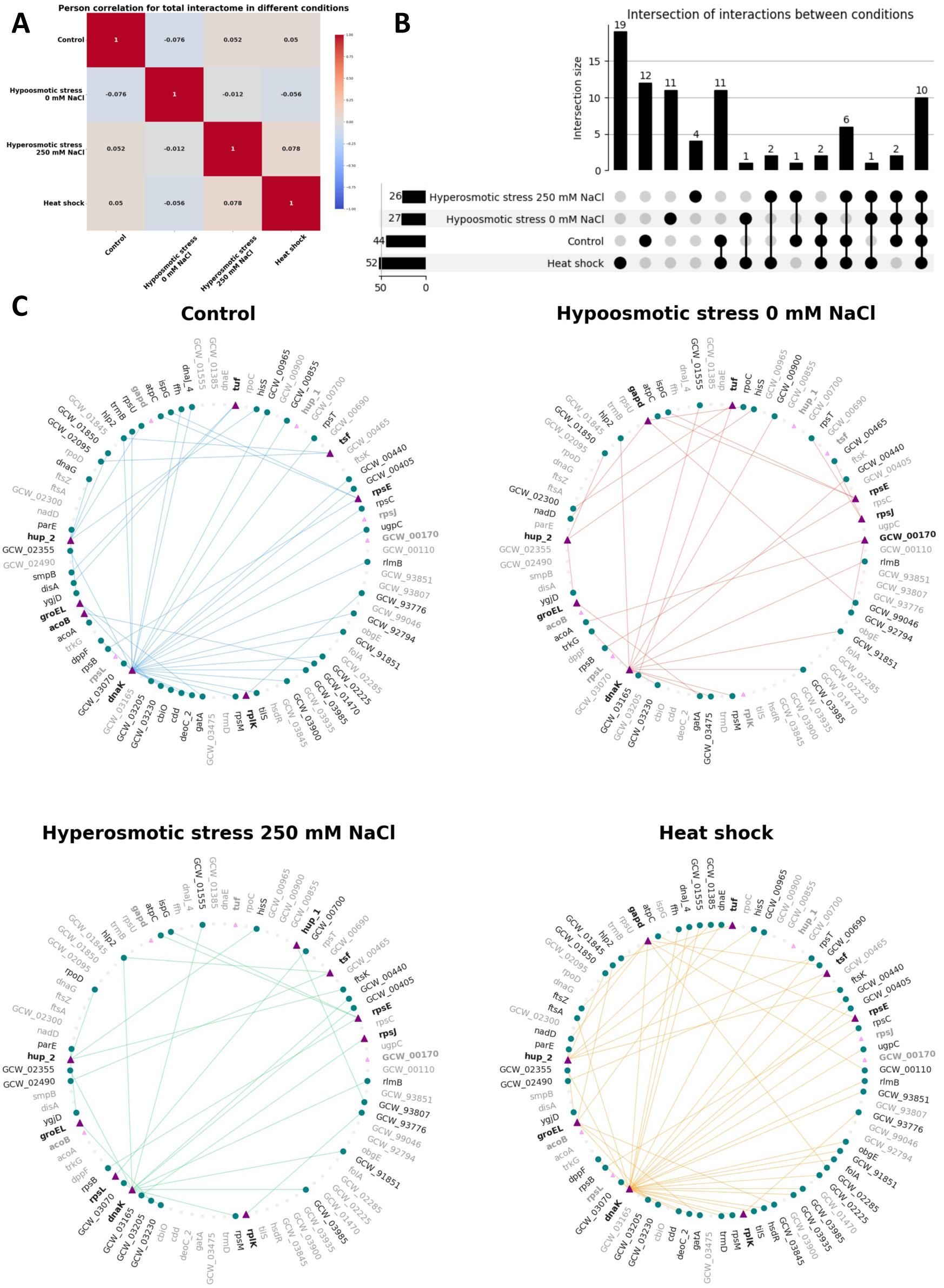
Characterization of the *M. gallisepticum* interactome under specific conditions. (A) Heatmap showing Pearson correlation coefficients for aggregate interaction frequencies across different stress conditions. (B) UpSet plot showing intersections between interactions involving Group 1 major proteins under each condition. (C) Interaction diagrams of major proteins under different stress conditions. Group 1 major proteins are shown as triangle nodes, partner proteins as circle nodes, and edges indicate interactions detected using the EGS cross-linker.

At the same time, the total number of interactions involving major proteins decreased under osmotic stress, whereas under heat shock it remained essentially unchanged (Figure 3c). Despite the large number of observed changes, the overall pattern of properties among the interaction partners of major proteins remained largely stable (Supplementary F1). However, differences from the overall proteome were detected: under physiological conditions, these partners were more soluble, had a higher positive charge, and showed higher functional temperature stability.

To test whether this type of protein organization is conserved across mycoplasmas, we analyzed published proteomic interactome data from *M. pneumoniae* cultures grown under normal conditions, obtained using the DSS cross-linking agent^52^. However, DSS differs from the EGS cross-linker used in our study, most notably in spacer arm length (11.4 Å for DSS versus 16.1 Å for EGS). We attribute possible discrepancies in the detected protein complexes to this difference. Nevertheless, for nearly one quarter of the *M. pneumoniae* proteins for which homologs in *M. gallisepticum* were identified by annotation (8/31), we observed matching complexation partners (Supplementary Table 4).

Furthermore, we analyzed whether the observed protein-protein interactions of Group 1 major proteins under different stress conditions were mediated by the same interaction interfaces. As shown in Figure 4 for the most interactive proteins, because Group 1 proteins are characterized by a high content of IDRs, different stress conditions appear to induce small conformational changes in these proteins, thereby altering the contact interfaces of their interaction partners.

**Figure 4.**
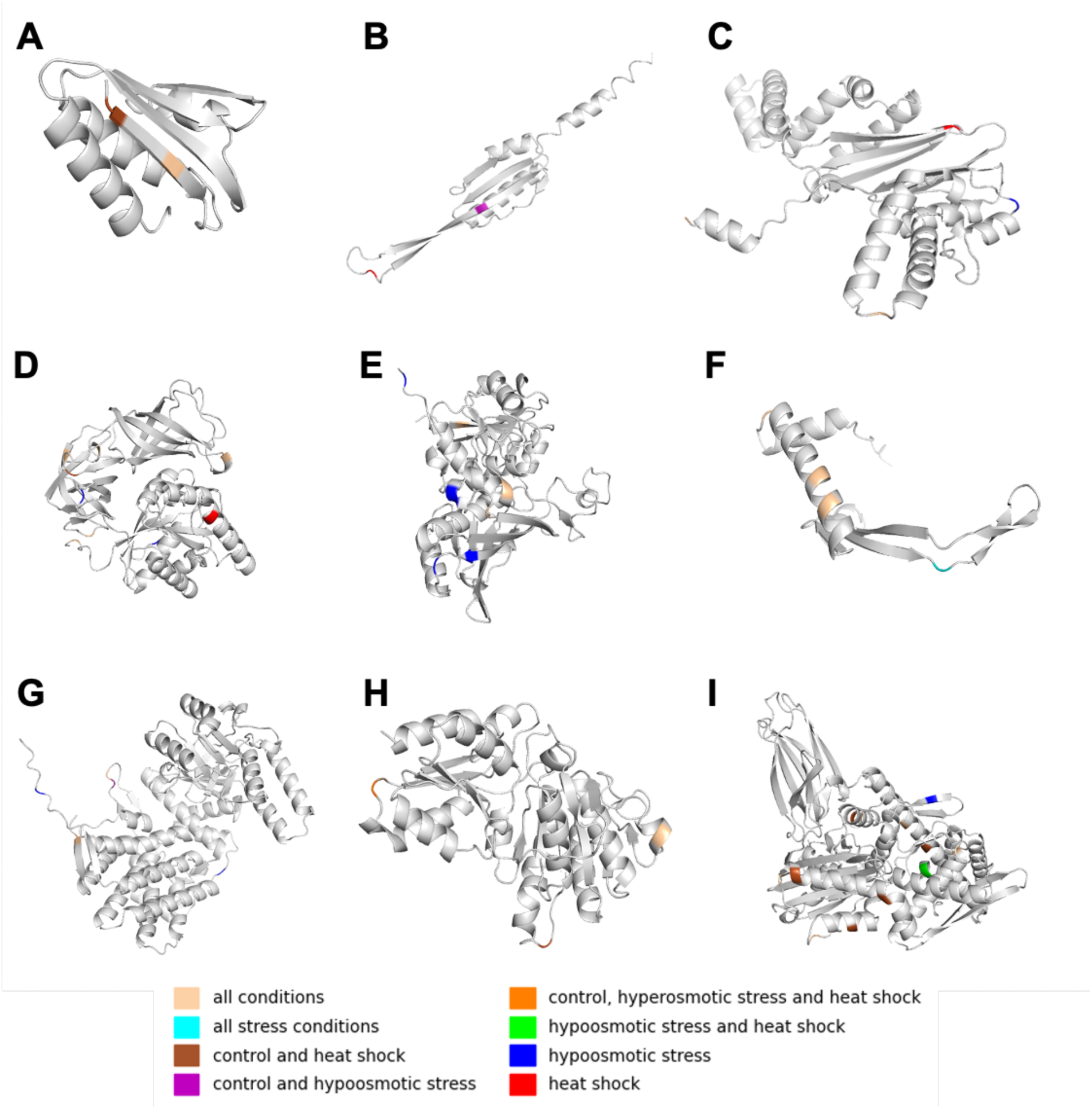
Binding sites of major *M. gallisepticum* proteins. Colored bars indicate conditions as shown in the figure legend. (A) RpsJ (A0A0F6CJV3). **(B)** Gapd (A0A0F6CKH9). **(C)** FruB (A0A0F6CJT4). **(D)** Tsf (A0A0F6CJY9). (E) Tuf (A0A0F6CKA9). (F) AcoB (A0A0F6CKX6). (G) Hup_2 (A0A0F6CKR5). (H) DnaK (A0A0F6CL29). (I) GroEL (A0A0F6CKW9).

### Minor Proteins in *M. gallisepticum* Are Intercorrelated

In minimal cells, not all proteins need to be equally abundant. Even in natural mycoplasmas and in the minimal bacterium JCVI-Syn3A, the abundance difference between major and minor proteins spans approximately 3.5 orders of magnitude. Protein abundance, therefore, cannot be equated with functional importance, since some proteins may be essential but toxic when overabundant^53^.

To examine the stability of protein abundance relationships, the concentrations of all proteins, both major and minor, were normalized in each sample using z-scores. Pearson correlation coefficients were then calculated for all protein pairs to identify highly coordinated abundance patterns across the dataset, which are shown as a heatmap in Figure 5a. The resulting matrix revealed distinct groups of highly correlated proteins, which were more clearly represented as graphs (Figure 5b). The data were filtered using correlation coefficient thresholds and visualized as weighted graphs, where vertices correspond to proteins and edges represent correlations above a given absolute threshold, with red indicating positive correlations and blue indicating negative correlations.

**Figure 5.**
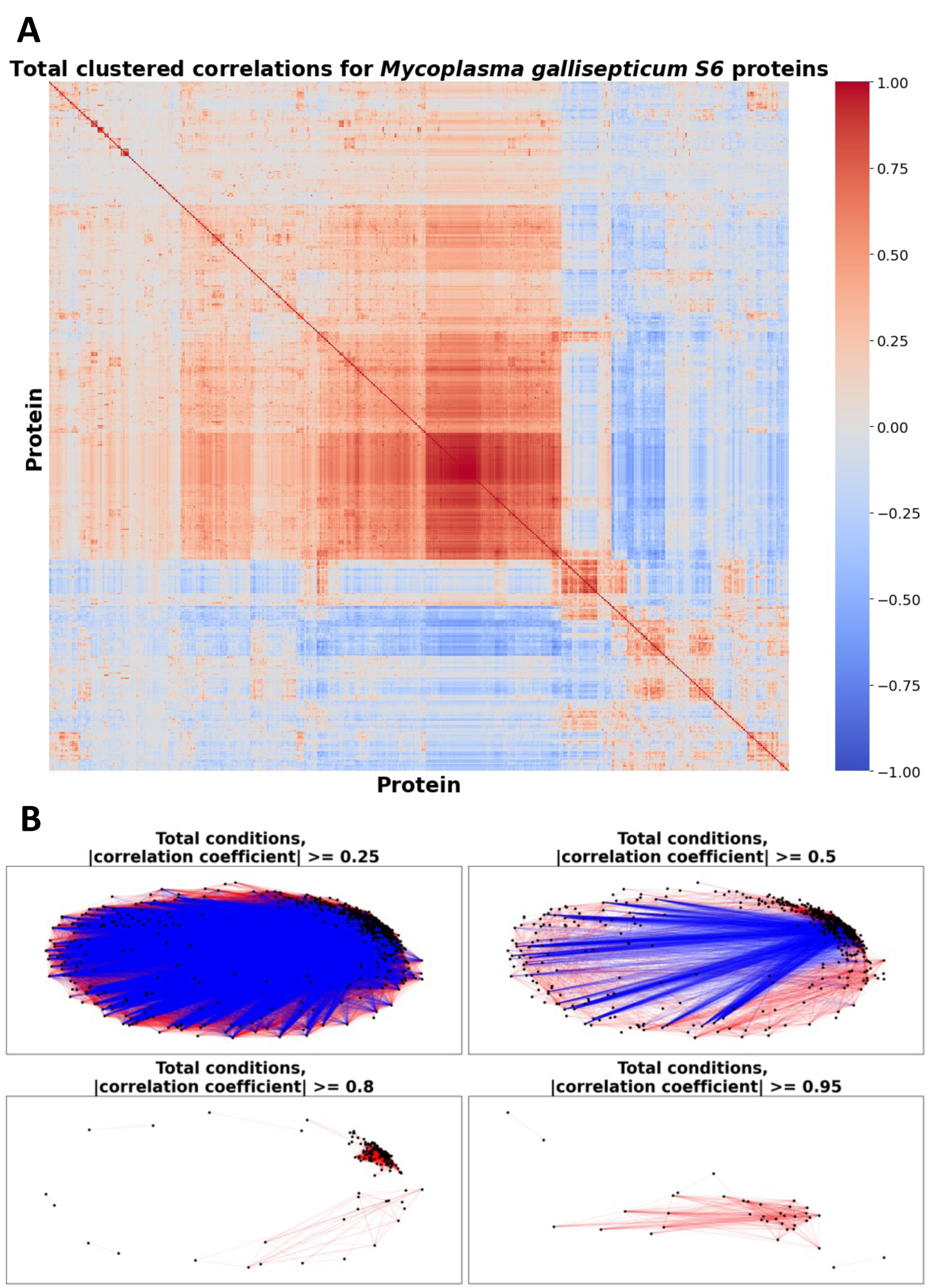
Protein-protein correlations in the *M. gallisepticum* proteome. (A) Heatmap showing Pearson correlation coefficients for quantitative protein-protein relationships across all conditions. (B) Graph representation of Pearson correlation coefficients for quantitative protein–protein relationships across all conditions. Vertices represent proteins, and edges represent correlations above a defined threshold, with red indicating positive correlations and blue indicating negative correlations.

At low correlation thresholds (≤0.75), both the heatmaps and graphs showed that the *M. gallisepticum* proteome separates into two opposing protein groups. One cluster was denser and persisted even at a threshold of 0.95, indicating exceptionally strong correlations among these proteins. To ensure that these patterns did not arise from peptide identification artifacts, peptide-level correlations within this group were also examined. Average inter-protein peptide correlations were higher than intra-protein peptide correlations, supporting the reliability of the observed protein-level relationships (Supplementary F2).

Analysis of the clustered proteins (Table 3) revealed no functional enrichment, except for an overrepresentation of proteins with unknown functions and enrichment for proteins containing transmembrane alpha-helices (17 out of 38). For 10 of these proteins, membrane integration (“integral component of membrane”) was confirmed in the Gene Ontology database^54^, and membrane transport functions were also noted. Among the remaining proteins, three ribosomal proteins, an acyl carrier protein, an endonuclease, a histidine-like protein, thioredoxin, and various enzymes distributed across metabolic pathways were identified. Consistent with their predominantly membrane localization, this group showed a shift in the GRAVY index toward hydrophobicity and a slight reduction in polar amino acid content.

**Table 3.** Proteins that have at least one partner with an absolute value of correlation coefficient ≥ 0.95.

| Protein category | Protein ID | Protein name | Gene | Gene |
| --- | --- | --- | --- | --- |
| **Ribosomal proteins** | A0A0F6CJW3 | Small ribosomal subunit protein uS17 | GCW_00325 | rpsQ |
|  | A0A0F6CLA9 | Small ribosomal subunit protein bS16 | GCW_03520 | rpsP |
|  | A0A0F6CL97 | Small ribosomal subunit protein bS18 | GCW_03455 | rpsR |
| **Nucleotides biosynthesis** | A0A0F6CKZ7 | Histidine triad nucleotide-binding protein | GCW_02870 |  |
|  | A0A0F6CL45 | DUF1600 domain-containing protein | GCW_03160 |  |
| **DNA-associated** | A0A0F6CK36 | Holliday junction resolvase RecU | GCW_00805 | recU |
|  | A0A0F6CL88 | Holliday junction branch migration complex subunit RuvA | GCW_03410 | ruvA |
| **tRNA ligases** | A0A0F6CK47 | Methionyl-tRNA synthetase | GCW_00860 | metG_1 |
| **VlhA** | A0A0F6CKK8 | VlhA.5.02 variable lipoprotein family protein | GCW_01970 |  |
|  | A0A0F6CKS3 | VlhA.3.06 variable lipoprotein family protein | GCW_02410 |  |
|  | A0A0F6CL79 | VlhA.3.03 variable lipoprotein family protein | GCW_03350 |  |
| **Adhesins** | A0A0F6CK79 | Uncharacterized protein | GCW_01095 |  |
| **Uncharacterized protein** | A0A0F6CK43 | Uncharacterized protein | GCW_00840 |  |
|  | A0A0F6CK13 | Uncharacterized protein | GCW_00650 |  |
|  | A0A0F6CKB3 | Uncharacterized protein | GCW_01355 |  |
|  | A0A0F6CKP9 | Uncharacterized protein | GCW_02220 |  |
|  | A0A0F6CLQ6 | Uncharacterized protein | GCW_03740 |  |
|  | A0A0F6CM36 | Uncharacterized protein | GCW_93849 |  |
|  | A0A0F6CL52 | Uncharacterized protein | GCW_03205 |  |
|  | A0A0F6CLJ0 | Uncharacterized protein | GCW_04035 |  |
| **Transporters** | A0A0F6CLT5 | Uncharacterized protein | GCW_90526 |  |
|  | A0A0F6CKY0 | Cation transporter | GCW_02765 | trkG |
|  | A0A0F6CKU9 | Permease | GCW_02595 |  |
| **Lipoproteins** | A0A0F6CL50 | Lipoprotein | GCW_03195 |  |
| **Other carbohydrate metabolic process** | A0A0F6CKM2 | ABC transporter ATP-binding protein | GCW_02060 |  |
|  | A0A0F6CLG9 | Aldehyde dehydrogenase | GCW_03880 |  |
| **Membrane protein** | A0A0F6CK07 | Membrane protein | GCW_00600 |  |
|  | A0A0F6CK56 | Membrane protein | GCW_00965 |  |
|  | A0A0F6CKT6 | Transmembrane protein | GCW_02515 |  |
|  | A0A0F6CL41 | Major facilitator superfamily permease | GCW_03140 |  |
|  | A0A0F6CKG9 | ATP synthase subunit a | GCW_01715 | atpB |
|  | A0A0F6CKN5 | RDD family protein | GCW_02150 |  |
|  | A0A0F6CLI8 | Phosphatidylglycerol-prolipoprotein diacylglycerol transferase | GCW_04020 | lgt |
|  | A0A0F6CKB1 | MATE efflux family protein | GCW_01330 |  |
| Others | A0A0F6CKA8 | Acyl carrier protein | GCW_01305 | acpP |
|  | A0A0F6CJZ3 | 1-deoxy-D-xylulose-5-phosphate reductoisomerase | GCW_00495 | dxr |
| Stress response | A0A0F6CL70 | Cytidine deaminase | GCW_03300 | cdd |
|  | A0A0F6CLB6 | Thioredoxin | GCW_03560 | trxA_2 |

\*These 38 proteins constitute the "constitutive cluster" described in the text

### Correlation Patterns Under Specific Conditions

Having observed strong correlations among a group of proteins across all conditions, the next step was to examine how the number of correlated proteins changes under individual stresses. Using the same correlation algorithm on smaller condition-specific datasets, the number of proteins with correlations above each threshold was plotted against the threshold value (Figure 6a). Correlations under control conditions closely mirrored the overall pattern, whereas specific stress conditions produced substantially more highly correlated proteins.

**Figure 6.**
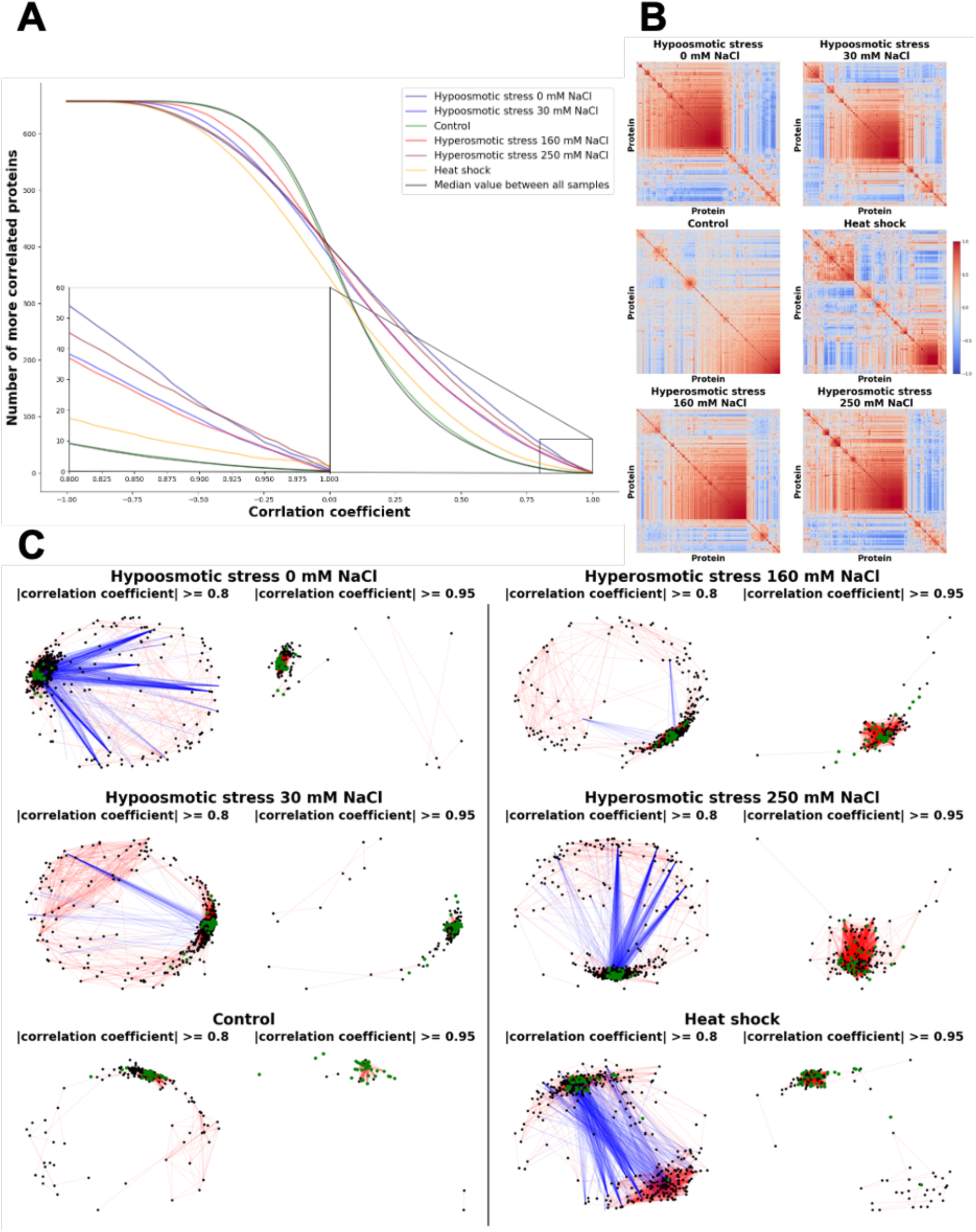
Protein-protein correlations in the *M. gallisepticum* proteome under specific conditionals. (A) Relationship between the correlation coefficient threshold and the number of proteins with at least one partner above that threshold. (B) Heatmap showing Pearson correlation coefficients between proteins analyzed separately for each condition. (C) Graph representation of Pearson correlation coefficients between proteins analyzed separately for each condition. Vertices represent proteins, with green vertices indicating the core of the constitutive cluster, and edges represent correlations above the threshold value, with red indicating positive correlations and blue indicating negative correlations.

Heatmaps and graphs were then constructed to explore these relationships in more detail (Figures 6b, c). The main cluster persisted across all conditions (green vertices), forming the core of a constitutive cluster that acquired additional proteins in each condition. In some conditions, alongside expansion of the constitutive cluster, a second cluster also persisted even at high correlation thresholds.

Regardless of condition, the constitutive cluster maintained consistent physicochemical property distributions while incorporating condition-specific proteins. It also retained enrichment exclusively for proteins with alpha-helical transmembrane domains (Supplementary Table 1). In contrast, the second, “variable” cluster persisted across all stresses at higher correlation thresholds and remained detectable even at the 0.95 threshold under hypoosmotic and heat shock conditions. This variable cluster showed no functional enrichment across most conditions (hypoosmotic stress 0 mM and heat shock) due to low diversity, except under hypoosmotic stress with 30 mM NaCl, where 3 of the 12 retained proteins were glycolytic enzymes (GpmL, PfkA, Fba) and 5 were ribosomal proteins (RplB, RplR, RplL, RpsK, RpmA). Physicochemically, proteins in these clusters differed significantly from the total proteome in size, charge, stability and reactivity, which may be related with their role in cellular adaptation (Supplementary F3).

## Discussion

Over more than 30 years of proteome research, technological capabilities have advanced dramatically. The field began with studies such as the work of Valerie C. Wasinger and colleagues, who used two-dimensional electrophoresis and MALDI-MS in the study of *Mycoplasmagenitalium*, where the term “proteome” was first introduced^55^. Today, proteomics enables large-scale measurement of thousands of cellular proteins in a single run using high-throughput, high-accuracy mass spectrometry^56,57^. Computational methods have advanced in parallel, enabling raw mass spectrometry data to be transformed into quantitative proteomic results^58^.

Despite these technological breakthroughs, a several major questions remain unresolved. These include how cellular regulatory mechanisms operate as a whole, how widespread epigenetic regulation is among bacteria, how epigenetic systems contribute to bacterial physiology and pathogenesis, and what role proteins play within these systems^59^. For example, recent evidence suggests that epigenetic regulation coupled with a long-term “memory” effect plays a major role in bacterial persistence formation^60^. Moreover, a recent study showed that soluble proteins inside a migrating cell do not move by random diffusion alone, but are instead carried by directed fluid flows^25^. These flows are generated by a specialized membraneless barrier composed of contractile molecules and act as a microscopic pump, delivering dissolved “building materials” directly to the leading edge of the cell.

Together, these findings further demonstrate that many regulatory and adaptive mechanisms in cells, both prokaryotic and eukaryotic, do not depend solely on the function of a specific gene or protein. Instead, they are also strongly shaped by the physicochemical properties of the cytoplasm. Alongside proteins, nucleic acids also contribute significantly to the spatial organization of the intracellular environment, primarily through liquid–liquid phase separation (LLPS) driven by electrostatic interactions. However, the molecular underpinnings of phase separation differ markedly between these two classes of biopolymers. In proteins, LLPS is typically governed by multivalent, weak, and transient interactions, including electrostatic and hydrophobic contacts, as well as contributions from intrinsically disordered regions (IDRs) and modular interaction domains. This diversity of interaction modes underpins the high selectivity and structural heterogeneity of proteinaceous condensates, which vary as a function of amino acid composition and the conformational plasticity of the participating proteins^61^.

In contrast, nucleic acid-driven phase separation relies predominantly on electrostatic associations between negatively charged phosphate backbones and cationic molecules or positively charged protein partners, often in coordination with complementary base-pairing interactions. Given the relative chemical uniformity of nucleotides compared to amino acids— particularly in terms of charge density and physicochemical versatility—the mechanisms governing nucleic acid condensation are inherently less specific. As a result, these processes are more strongly influenced by polymer length, secondary structure, and local concentration than by the broad spectrum of interaction chemistries that characterize protein-based condensates.

On the regulatory side, the ability of cells to modulate local mRNA concentrations independently of total protein abundance appears to play a key role in reorganizing the density and compositional landscape of ribonucleoprotein (RNP) condensates. Given that the global mRNA pool remains largely constant, this phenomenon likely reflects a spatial redistribution of transcripts across subcellular compartments rather than global changes in gene expression or *de novo* aggregation events. Such redistribution may serve as a rapid, post-transcriptional mechanism for tuning the material properties and functional output of RNP granules in response to cellular cues. However, given the high protein-to-mRNA ratio in the cell, the quantitative and physicochemical properties of proteins are likely to dominate cytoplasmic organization, making their systematic characterization essential for understanding the regulatory architecture of the cell.

Cellular regulation is distributed across multiple levels, with proteins acting not only as enzymes, but also as signals and structural components. A single protein can therefore operate as a node in several networks at once. As a result, a regulatory “mechanism” is rarely confined to one pathway, and defining only the individual function of a protein is not enough to explain its role in the cell. Regulatory networks may also be nonlinear and redundant, making the system resilient to perturbations and difficult to reconstruct causally^62,63^.

Bacteria of the genus *Mycoplasma* provide a useful model for exploring these principles. Their naturally reduced genomes make them minimal cells, yet previous studies have shown that they maintain remarkable proteome stability despite external and internal perturbations^5,6,64^. This is particularly intriguing because mycoplasmas, unlike more complex bacteria such as *E. coli* and *B. subtilis*, possess relatively few transcriptional and translational regulatory factors^11^. We therefore hypothesized that these minimal cells may rely on more general physical mechanisms for regulation and adaptation.

Despite this proteome stability, mycoplasma cells clearly adapt to changing environmental conditions, as reflected by metabolic changes^64^ and broad resilience across a wide range of stresses. We therefore analyzed proteome dynamics as an integrated whole in *Mycoplasma gallisepticum* under various conditions.

First, we performed quality validation of mass spectrometry data accumulated by our group on *M. gallisepticum* proteomes under various perturbation, including control conditions, heat shock, and osmotic stresses. From this dataset, we selected 504 samples with high technical quality and maximal procedural uniformity in both sample preparation and analysis (Figures 1a-c).

These curated datasets were then used to determine how rapidly the total *M. gallisepticum* proteome saturates with protein mass (Figure 1d). Based on these calculations, we defined the major protein core of the cell, consisting of Group 1 and Group 2 proteins, which comprised 17 and 181 proteins, respectively. We then compared the functional composition of the major and minor protein groups (Figure 1e). Despite differences in the proportions of individual functional categories between the major and minor protein fractions, both groups remained highly heterogeneous and covered all annotated categories.

Under stress conditions that affect cellular metabolic rate, the contribution of these major proteins to cellular biomass increased. We hypothesize that this shift may influence cytoplasmic organization, protein condensation processes, and the range of cellular reactions, since the physicochemical properties of the most abundant proteins (Group 1) differ from those of the rest of the proteome (Figure 2a).

Group 1 proteins are distinguished by several shared physicochemical features. They show a greater positive charge, largely due to the abundance of histone-like and ribosomal proteins, reduced average hydrophilicity reflected by a lower GRAVY index, and low temperature sensitivity, which may make their functional and structural activity more stable under external influences. Furthermore, these proteins are classified as nucleic acid-binding proteins, reflecting their importance and potential involvement in the regulation of genomic information. Moreover, the transcription of genes encoding two major ribosomal proteins, RpsJ and RpsE, is regulated by WhiA, one of the limited number of transcriptional regulators encoded in the mycoplasma genome, a finding that may underscore the physiological importance of maintaining stable intracellular concentrations of these proteins^14^.

These proteins also contain more potential IDRs and more potential reactogenicity sites, which, under conditions of high protein concentration, may correlate with a scaffold-like function^65^. This possibility is consistent with several known examples: Gapd is involved in “metabolon” complex formation for the regulation of biocatalysis^66^, chaperones and histone-like proteins cooperate to establish condensed DNA assemblies^67^, and ribosomal proteins have also been linked to phase formation^68^.

Given that IDR-containing proteins can induce LLPS, Group 1 major proteins may serve as suitable components for the dynamic formation of biomolecular condensates and membraneless compartments capable of supporting temporal regulation inside the cell. Structural disorder is therefore not merely a secondary feature of proteins, but an important functional property, even in minimal organisms such as mycoplasmas^69^. Moreover, most of the identified IDRs were also found in other organisms (Supplementary Table 2), suggesting that they may play a broader role in organizing the prokaryotic cytoplasm.

At the functional level, protein-mediated phase separation can be modulated by extrinsic physical stimuli. For instance, mechanical deformation of the cell wall induces reorganization of membrane-associated and submembrane proteins, leading to redistribution of surface charges and enhanced cellular adhesion to specific substrates^70^. Furthermore, external electric fields exert pronounced effects on bacterial physiology, influencing viability, behavior, and metabolic state through alterations in membrane potential and permeability, as well as by modulating motility, growth kinetics, and overall functional activity^71^.

In addition, evidence supports that Group 1 proteins constitute the physical base of minimal core of the cell and can exert a significant influence on cellular state not only through their abundance, but also through their role as a universal attractor. This is supported by the observation that most proteins in this group, excluding ribosomal and mycoplasma-specific proteins, are included in the universal minimal interactome^72^.

Comparison of these proteins with the proteomic composition of minimal cell models, JCVI-Syn3A and *M. gallisepticum* nanoforms, revealed substantial overlap across the three groups (Figure 2b). The overlapping proteins covered key cellular processes essential for viability (Figure 2c), supporting the use of this approach to define the minimal protein core required for mycoplasma cell viability.

Correlating protein abundance with functional importance remains challenging. Some proteins may be essential yet toxic when overabundant, whereas others may perform organizing or moonlighting functions simply because of their high abundance. We therefore investigated whether *M. gallisepticum* proteins exhibit coordinated behavior independent of stress conditions. This analysis revealed a group of 38 low-abundance proteins, referred to here as the “constitutive cluster,” that displayed remarkably coordinated abundance changes. These changes were globally unrelated to known co-expression patterns, operon localization, or complex formation (Figure 5, Table 3). Despite their low abundance, proteins in this cluster participate in several key cellular processes, including translation, lipid biosynthesis, DNA recombination and repair, and transmembrane transport of various substances.

This effect became more pronounced when the analysis was restricted to individual conditions. Under stress, the *M. gallisepticum* proteome showed not only co-clustering of the conserved core proteins with condition-specific partners, but also the formation of a second, variable correlation cluster under some conditions (Figure 6). Notably, these clusters did not simply coexist; they were anticorrelated, suggesting a possible role in balancing and stabilizing the proteome under stress.

However, these relationships could not be confirmed at the interactome level. This may reflect the low abundance of the interacting partners or specific features of their interactions, such as different inter-partner distances, alternative interaction mechanisms, or mediation through a bridging protein.

Likewise, in the interactome obtained using the EGS cross-linker, we observed a global rearrangement of interactions involving core proteins (Group 1) across different stress conditions (Figure 3a). This rearrangement may be related to regulation of the cellular response to stress. The variation in partners identified for these core proteins may, in part, reflect the increased content of unfolded regions within them.

Under different stress conditions, the conformation of a protein may be slightly modified through its IDRs, leading to the formation of new transient protein pairs. This is partly supported by the observation that, for the same Group 1 protein, different binding peptides were detected with its partner protein under different stress conditions, including peptides located in unfolded protein regions (Figure 4).

This may also explain why no significant changes in *M. gallisepticum* protein abundance are observed under stress conditions, despite the bacteria’s apparent adaptation. Rather than changing overall protein abundance, the cell appears to respond by rearranging proteins within complexes, which may represent a more energy-efficient strategy. Such dynamic variability of protein states, coupled with quantitative stability, may allow *M. gallisepticum* to achieve high adaptive capacity. At the same time, 10 interaction patterns involving Group 1 major proteins were preserved across all conditions (Figure 3b). These included, for example, interactions of RpsE with AtpC and RpsB, which have been reported in the literature as predictors of the bacterial cell cycle^73^ and as ribosomal neighbors, respectively. DnaK showed the greatest number of stable interactions, consistent with its active involvement in the stress response^74^, including through interactions with various sigma factors, one of which was also detected in our results.

Interestingly, only osmotic stresses led to a significant change in the total number of partners interacting with major proteins (Figure 3c). This may first be related to the substantial changes in salinity experienced by the bacterium and, consequently, to changes in the protein environment. Second, these stresses are chronic: mycoplasma cells were grown at these saline concentrations until the mid-logarithmic growth phase. Thus, the observed rearrangement of protein interactions could reflect adaptation to prolonged stress.

Under heat shock, restructuring of protein complexes was also observed, although the overlap with the control condition was greater than under osmotic stress (Figure 3c). This may be related to differences in intracellular metabolite concentrations between stress and normal conditions. In our previous studies, we showed that ATP concentration increases in *M. gallisepticum* cells under stress conditions^64^. Since ATP at high concentrations can also function as a hydrotope^33^, stress conditions may alter the solubility, and consequently the properties, of mycoplasma proteins. This, in turn, could be associated with their rearrangement into transient complexes.

Despite these changes, the property distributions of the major proteins’ partners remained highly similar across conditions (Supplementary F1). This may reflect underlying principles that shape the regulation and organization of the cellular response. Moreover, the composition of the soft condensed phase is known to influence cytoplasmic organization^75^, which may also contribute to the cellular response to stress.

Furthermore, the observed protein interactions are not unique to *M. gallisepticum*. We analyzed published interactome data for *M. pneumoniae* obtained using the DSS cross-linking agent^52^. Although the spacer length of DSS differs from that of the EGS agent used in our work, which may affect the identification of protein partners, nearly one quarter of the proteins with corresponding annotated homologs between the two mycoplasma species shared common protein– protein interactions (Supplementary Table 4).

We hypothesize that, in many cases, the formation of a rigid protein–protein pair may not be strictly necessary. Instead, similarity in the physicochemical properties of these proteins may be sufficient to ensure that the space they occupy is properly “structured.” Thus, the observed overlaps may point to a universal regulatory mechanism in mycoplasmas, mediated by the formation of such transient protein complexes. These include canonical complexes as well, such as DnaJ-DnaK^76^, Ffh-DnaK^77^ and AcoA-AcoB^78^ complexes.

As highlighted above, the physicochemical properties of the cytoplasm—shaped by both proteins and nucleic acids—appear to play a central role in these adaptive responses. The ability of Group 1 proteins to undergo LLPS via IDRs, coupled with the nucleic acid-driven spatial redistribution of transcripts and the modulation of condensate density, suggests that *M. gallisepticum* may leverage broad biophysical principles to compensate for its limited regulatory repertoire.

In conclusion, our study demonstrates that quantitative stability of the *M. gallisepticum* proteome does not imply absence of proteome-level stress responses. Rather, in these bacterial cells, adaptation may occur through changes in proteome organization that are not reflected by major shifts in protein abundance. Possibly because of their substantial genome reduction, mycoplasmas may rely on more primitive mechanisms of adaptation and regulation that do not depend on specific regulators, but are instead governed by broader physical principles.

We found that *M. gallisepticum* exhibits dynamic proteome behavior as an integrated whole. A stable core of major proteins was identified, which appears to shape the overall physicochemical state of the cytoplasm. These proteins contain a high proportion of IDRs, which may contribute both to LLPS and to the formation of distinct protein complexes in response to different adaptive challenges. Minor proteins, in turn, form one or two intercorrelated clusters depending on external conditions.

Together, these findings suggest that *M. gallisepticum* may adapt not by substantially changing the abundance of its proteins, but by reorganizing proteome architecture and rearranging protein complexes. Such a strategy could allow the cell to conserve limited energy resources while maintaining high adaptive capacity within cell populations, supporting survival under diverse stress conditions.

## Material and methods

### *Mycoplasma gallisepticum* cell culture and stress conditions

*M. gallisepticum* S6 cells were cultivated, and nanoforms were obtained as described previously^49^. Osmotic stress conditions were generated by adjusting the NaCl concentration in the cultivation medium. Hypoosmotic conditions were achieved either by omitting NaCl from the medium or by reducing its concentration to 30 mM, although trace amounts remained due to the presence of yeast dialysate and horse serum. Hyperosmotic conditions were established by increasing the NaCl concentration to 160 or 250 mM, compared with the standard concentration of 85 mM. Heat shock was induced by incubating cells at 46 °C for 30 min.

### *In vivo* cross-linking

*M. gallisepticum* S6 pellets were washed twice in PBS to remove amine-containing components from the medium. The cells were then resuspended in PBS at ∼10⁹ cells/mL and cross-linked by adding ethylene glycolbis(succinimidylsuccinate) (EGS) (Thermo Scientific, USA) to a final concentration of 5 mM for 30 min at room temperature. The reaction was quenched by adding Tris-HCl, pH 8.0, to a final concentration of 20 mM for 15 min at room temperature. Cells were collected by centrifugation at 8000 × g for 10 min at 4 °C.

### Proteomic analysis

Sample preparation, comprising protein extraction and proteolytic digestion, as well as LC– MS/MS analysis of the resulting peptide samples, was carried out in accordance with a previously described protocol^24^.

### Data analysis

Quantitative analysis of MS/MS data was performed using MaxQuant software^79^, with peptide identification carried out by the Andromeda search engine against the *Mycoplasma gallisepticum* S6 genome annotation database. For each sample, relative protein abundance was calculated as the ratio of the protein’s intensity-based absolute quantification (iBAQ) value to the summed iBAQ intensity of all detected mycoplasma proteins in that sample. Within each experimental group, median relative intensities across biological replicates were calculated, and fold changes were determined relative to the control condition.

For cross-linking samples, raw files were processed into peak lists using ProteoWizard (version 3.0)^80^. Cross-linking MS/MS data were then analyzed using xiSEARCH^81^. The search parameters were as follows: MS accuracy, 6 ppm; MS/MS accuracy, 20 ppm; enzyme, trypsin; cross-linker, EGS; maximum missed cleavages, 5; missing monoisotopic peaks, 2; fixed modification, carbamidomethylation on cysteine; variable modifications, oxidation on methionine and deamidation on asparagine and glutamine; and fragments, b and y ions with loss of H₂O, NH₃, and CH₃SOH.

For each sample, the presence or absence of each interaction was determined. Interactions detected in two out of three technical replicates were considered statistically significant and assigned to the corresponding biological replicate. Interaction frequency was then defined as the ratio of biological replicates in which the interaction was detected to the total number of biological replicates.

### Protein property calculation

Protein structures were predicted using AlphaFold^41^. Protein volumes and packing densities were then calculated using the Java-based ProteinVolume application^40^. For each structure, the Energy Minimize parameter was set to true, be because hydrogen atoms are absent from AlphaFold-predicted structures.

The ExPASy ProtParam module was used to calculate the isoelectric point, total number of charged residues, GRAVY, instability index, and aliphatic index^39^. Intrinsic disorder propensity was predicted for each amino acid using IUPred2A, after which the fraction of amino acids with a disorder score of at least 0.5 was calculated^42^.

Reactogenicity was predicted using ISPRED-SEQ^43^, which estimates the probability of each amino acid interacting with other proteins. For each protein, the maximum amino acid value was recorded. To estimate the number of potential reactogenicity centers, the proteome-wide median reactogenicity value was used as a threshold. The number of amino acids in each protein with an interaction probability above this threshold was then calculated.

Solubility was predicted using the CamSol web server, with the pH parameter set to 7^44^. Dipole moments were calculated using the ProteinDipole script^45^. Functional stability temperature was predicted using TemStaPro software in advanced mode with additional temperature thresholds^46^. Melting temperature was predicted using DeepSTABp^47^, with the growth temperature set to 37 °C and the melting environment set to “cell.”

For major proteins, DNA-binding probability was estimated from AlphaFold-predicted structures using the DP-Bind web-server^48^. Since DP-Bind predicts a value for each amino acid, the fraction of amino acids predicted to bind DNA was used for subsequent calculations.

When constructing the distributions, the mass contribution of each protein to the average values across all conditions was taken into account. For Supplementary Figure 3, mass contributions from the corresponding stress conditions were used.

### Multiple protein alignment

Multiple sequence alignment of protein sequences was performed using Clustal Omega^82^. Amino acid sequences for *M. gallisepticum S6*, *M. genitalium*, *M. pneumoniae*, the synthetic bacterium JCVI-Syn3A, *E. coli*, *B. subtilis*, *Pseudomonas aeruginosa* and *Staphylococcus aureus* were downloaded from UniProt^83^ in FASTA format. Alignments were performed using default settings. The resulting multiple alignments were used to analyze amino acid residue conservation and to compare intrinsically disordered regions within the proteins.

## Statistical Analysis

The Pearson correlation coefficient (r) was used to assess the strength and direction of the linear relationship between variables. The statistical significance of each correlation coefficient was evaluated using a two-tailed t-test, with correction for multiple comparisons applied using the Benjamini–Hochberg false discovery rate (FDR) procedure. A result was considered statistically significant if the corresponding adjusted p-value was less than 0.01.

## Data availability

The raw proteomic data generated in this study have been deposited in the PRIDE database under the accession numbers PXD078057 for proteomic data and PXD077082 for interactomic data.

## ACKNOWLEDGMENT

The work was funded by Federal Service for Surveillance on Consumer Right Protection and Human Wellbeing, grant No. 122030900107-3.

## ASSOCIATED CONTENT

**Supplementary figure 1-3**. Physicochemical characterization of protein groups. (PDF)

**Supplementary table 1.** Characterization of *M. gallisepticum* protein. (XLSX)

**Supplementary table 2.** Sequence alignment results of major proteins for model organisms and predictions of disordered regions. (XLSX)

**Supplementary table 3.** Comparison of JCVI-Syn 3A and *M. gallisepticum* proteins. (XLSX)

**Supplementary table 4.** *M. gallisepticum* interactome obtained using EGS cross-linker. (XLSX)

